# Development of an optimized machine learning approach for assessing brain metastatic burden in preclinical models

**DOI:** 10.1101/2024.08.21.608131

**Authors:** Jessica Rappaport, Quanyi Chen, Tomi McGuire, Amélie Daugherty-Lopès, Romina Goldszmid

## Abstract

Brain metastases (BrM) occur when malignant cells spread from a primary tumor located in other parts of the body to the brain. BrM is a deadly complication for cancer patients and currently lacks effective therapies. Due to the limited access to patient samples, preclinical models remain a valuable tool for studying metastasis development, progression, and response to therapy. Thus, reliable methods for quantifying metastatic burden in these models are crucial. Here, we describe step by step a new semi-automatic machine-learning approach to quantify metastatic burden on mouse whole-brain stereomicroscope images while preserving tissue integrity. This protocol utilizes the open-source, user-friendly image analysis software QuPath. The method is fast, reproducible, unbiased, and provides access to data points not always obtainable with other existing strategies.

## 1 Introduction

Brain metastases (BrM) are an increasingly common and deadly complication in cancer patients (Achrol et al., 2019; Ahmad et al., 2023; Khan et al., 2023). Although various primary tumor types metastasize to the brain, lung cancer, breast cancer, and melanoma, are the three most frequent tumors to do so (Achrol et al., 2019; Khan et al., 2023; Strickland et al., 2022). Importantly, 40-60% of patients with metastatic melanoma develop BrM (Eroglu et al., 2019; Internò et al., 2023). Furthermore, among adult brain cancer patients, the incidence of BrM is higher than that of primary brain tumors (Khan et al., 2023; Lah et al., 2020; Lee et al., 2022; Rios-Hoyo and Arriola, 2023). The standard of care for this disease consists of surgical resection, whole-brain radiotherapy, and/or stereotactic radiosurgery, however, these are not effective for many patients and often result in tumor recurrence and various side effects (Achrol et al., 2019; Ahmad et al., 2023; Mut, 2012; Suh et al., 2020). While immune checkpoint blockade and targeted therapies have shown promising results in recent clinical trials, they benefit only a limited subset of BrM patients (Ahmad et al., 2023; Brastianos et al., 2023; Jänne et al., 2022; Long et al., 2018; Reungwetwattana et al., 2018; Rios-Hoyo and Arriola, 2023; Tawbi et al., 2021). Thus, new therapeutic strategies are needed to improve BrM outcomes.

Access to patient samples is limited, therefore, preclinical models remain critical tools to study BrM and treatment options. The main methods used for quantifying metastatic burden in preclinical models include in vivo/ex vivo imaging such as bioluminescence when labeled tumor cells are used, manual counting if lesions are visible, and histology (Miarka and Valiente, 2021; Valiente et al., 2020; Wu et al., 2020). Each of these methods have their advantages and disadvantages (Table 1) (Ansari et al., 2016; Day et al., 2022; Genevois et al., 2016; Klerk et al., 2007; Valiente et al., 2020). In vivo imaging for example, allows for longitudinal tumor burden assessment. However, the addition of foreign proteins (e.g., luciferase) renders tumor cells more immunogenic, which poses a problem when performing therapeutic studies (Ansari et al., 2016; Day et al., 2022; Grzelak et al., 2022). Other drawbacks of this approach are the difficulty in determining whether the signal originates from only one or several lesions, the risk of false negatives when lesions are too small and the signal does not surpass background levels, or when signal quenching occurs due to animal fur, hemoglobin, melanin from pigmented skin or tumor cell lines (Klerk et al., 2007; Valiente et al., 2020). Manual counting is inexpensive and does not require special equipment, but it is time-consuming and introduces operator bias (Bankhead et al., 2017; Lopès et al., 2018; Pennig et al., 2021). In addition, accurately measuring individual metastatic foci is very challenging with this technique. Histology provides unambiguous pathological assessment and allows for molecular characterization (Valiente et al., 2020; Wu et al., 2020). With this technique, one can also determine lesion size and total metastatic area. However, several serial sections from different tissue depths are needed to accurately quantify metastatic burden, which increases analysis time (Valiente et al., 2020; Wu et al., 2020). Moreover, because of the way samples are processed, they cannot be used for other types of analyses (e.g., high-parametric immune profiling).

**Table 1.** Advantages and limitations of different methods to quantify metastatic burden.

| **Approach** | **Advantages** | **Limitations** |
| --- | --- | --- |
| In Vivo Bioluminescence Imaging | • Longitudinal studies possible<br>• Quantification of metastases not visible without a reporter system | • Undesirable impact on tumor cell and host (e.g., immunogenicity)<br>• No detection of small metastases<br>• Possible signal quenching (e.g. animal fur, melanin)<br>• Individual metastases count not accessible or inaccurate<br>• Individual metastases size not accessible or inaccurate<br>• Spatial localization imprecise |
| Ex Vivo Bioluminescence Imaging | • Quantification of metastases not visible without a reporter system<br>• Spatial localization accessible<br>• Overcomes in vivo signal quenching | • All as above except signal quenching<br>• Longitudinal studies not possible<br>Tissue cannot be re-used to look at other parameters (e.g.: immune microenvironment by flow cytometry) |
| Macroscopic manual counting (on a whole organ using a microscope) | • Metastases count accessible<br>• Macroscopic spatial localization accessible<br>• Macroscopic morphological features accessible | • Biased from person-person<br>• Metastases size not accessible<br>• Time-consuming<br>• Longitudinal studies not possible |
| Histology | • Metastases count accessible<br>• Microscopic spatial localization accessible<br>• Microscopic morphological features accessible<br>• Limited cellular/molecular characterization possible | • Tissue cannot be re-used to assess other parameters<br>• Limited to the area on the slide being analyzed<br>• Time-consuming<br>• Longitudinal studies not possible |
| Our approach (e.g.: macroscopic metastasis burden assessment using a machine learning algorithm) | • Metastases count accessible and accurate<br>• Metastases total area and individual size are accessible and accurate<br>• Tissue can be re-used to assess other parameters<br>• Macroscopic morphological features accessible<br>• Macroscopic spatial localization accessible<br>• No artificial impact on tumor cell and host<br>• Unbiased | • Metastatic lesions need to be visible. (See note 22)<br>• Longitudinal studies not possible<br>• Optimal for assessment of superficial metastases. Alternative can be found to assess deeper lesions (See note 23). |

Here, we developed a semi-automatic machine-learning approach to quantify metastatic burden on whole-brain stereomicroscope images using the open-source, user-friendly software QuPath (Bankhead et al., 2017). This software was initially designed to perform digital pathology analysis on entire microscopic tissue slides. It allows the detection and annotation of different regions and/or cells of interest using machine learning techniques and an “object-oriented” program. Therefore, the user can create their own set of rules and workflow tailored to their specific scientific query. For instance, to automatically detect a particular area, the first step involves creating an “object” by selecting the structure of interest (e.g., tumor area) using QuPath’s drawing or automatic segmentation functions within a part of the image. Next, this newly created “object” can be annotated under a user-defined class (e.g., tumor) using different classifiers based on selected machine learning algorithms (e.g., Random Tree, Artificial Neural Network) and image properties (e.g., pixel values, neighborhood objects). In the same manner, multiple classes of objects are generated on different representative areas of the image, allowing the program to distinguish each region of interest. This process constitutes the machine learning “training” steps. These classes are then automatically applied by the program to the entire image during the “prediction” steps, allowing the annotation of all the structures present in the whole slide (Bankhead et al., 2017; Greener et al., 2022). Such an approach is commonly used by various software to analyze bioimages from microscopic slides, providing a powerful tool for measuring microscopic areas and quantifying various cell types in particular tissue compartments (Bankhead et al., 2017; Lopès et al., 2018; Maric et al., 2021). Its advantage lies in allowing unbiased, accurate, and quantitative biological analysis on whole-scanned slides in a timely and reproducible manner (Anwer et al., 2023; Bankhead et al., 2017; Greener et al., 2022). However, its use on a whole bright-field macroscopic organ image remains challenging, partly due to the image properties such as the high heterogeneity of tissue brightness induced by the 3-dimensional structure of the organ itself.

In this report, we provide a detailed protocol for optimizing the acquisition of macroscopic organ images and assessing brain metastatic burden using a newly developed workflow and adapted classifiers from QuPath (Bankhead et al., 2017). This machine learning-based method, which preserves tissue integrity, overcomes the limitations of other existing approaches (Table 1). Additionally, we offer technical and computational guidance for applying this method to other models and with different imaging equipment.

## 2 Materials

### 2.1 Common Disposables

- Transparent 6-well plate (Costar, catalog: 3056) (see 1)
- 20mL syringe (BD, catalog: 302830) (see 1)
- 21G needle (BD, catalog: 305165) (see 1)
- Bench pad
- Ice
- 6cm Petri dish (Thermo Fisher, catalog: 150288) (see 1)
- 3ml Transfer pipette (Falcon, catalog: 357575) (see 1)
- 15mL falcon tubes (if fixing the brain after imaging) (Corning, catalog: 430152) (see 1)
- Colored post-it (see 2)

### 2.2 Reagents

- Cold 1X Phosphate-Buffered Saline (PBS) (Corning, catalog: 21-040-cv) (see 1)
- Cold 1X PBS + 2mM EDTA
- 70% Ethanol prepared in PBS
- Cold RPMI (Thermo Fisher, catalog: 11875119) (see 1)
- Appropriate reagent for use or storage of the brain after imaging (see 3)

### 2.3 Equipment

- Curved forceps for brain harvest (Fisher, catalog: 12-460-518) (see 4)
- Blunt forceps for imaging (Fisher, catalog: 10-000-469) (see 4)
- Scissors (curved) (Fisher, catalog: 08-935) (see 1 and *4*)
- Leica M165 FC Stereomicroscope (see 5)

- Zoom range: 7.3x – 120x
- Resolution: max 453 lp/mm
- Working distance: 61.5 mm
- Magnification: 960x
- Object field: 63 mm
- Ring light attachment for microscope (Leica, LED5000 RL) (see 1 and 6)

### 2.4 Software

- LAS X (Leica Image Application Suite) (see 7)
- QuPath (Bankhead et al., 2017)
- Microsoft Excel
- GraphPad Prism 10

## 3 Methods

### 3.1 Material preparation

1. Generate in vivo mouse brain metastasis by implanting the desired tumor model in accordance with institutional Animal Care and Use Committee (ACUC) guidelines. Any model with lesions that can be distinguished from normal tissue can be used (e.g., intracarotid injection of 2×10^6^ cell/mL K1735 melanoma cells). (see 8)
2. The following materials can be prepared the day before sample collection:

a. 6-well plates with 5mL RPMI per well, store overnight at 4ºC
b. 1X PBS + 2mM EDTA (20mL/mouse), store overnight at 4ºC

### 3.2 Sample collection

1. Set up working area with bench pad. Keep all reagents on ice.
2. Fill syringe with 20mL PBS + 2mM EDTA, 1 syringe per mouse and keep on ice until needed for heart perfusion (see 9).
3. Euthanize metastasis-bearing mouse (see 10) in accordance with institutional ACUC guidelines.
4. Place mouse in dorsal recumbency and spray the ventral region with 70% ethanol.
5. Using scissors, make a midline incision through skin and peritoneal membrane all the way up to the xyphoid. Use forceps to carefully pull skin up to avoid organ damage.
6. Using scissors, cut diaphragm along the rib cage.
7. Use scissors to make two cuts through both sides of the rib cage. Using curved forceps, gently lift the sternum up or cut it off to fully expose the heart and lung to prepare the area for heart perfusion as previously described (Wu et al., 2021), (see 11 and 12).
8. Use curved forceps to move small intestines out of the body cavity to reveal the inferior vena cava. Cut the inferior vena cava to allow blood draining from tissues (see 11)
9. Insert the 21G needle from the syringe filled with PBS/EDTA halfway into the left ventricle and inject 10mL of PBS/EDTA with steady pressure. The liver should become bloodless and look pale (see 11).
10. Insert the needle into the right ventricle of the heart and inject the leftover 10mL PBS/EDTA with steady pressure. All lung lobes should become bloodless as evidenced by a whitish color (see 11).
11. Place the mouse body in a ventral position and remove the skin over the head to reveal the skull. Use scissors to cut the muscle between the neck and the base of the skull to reveal the occipital bone (Figure 1A). Expose the foramen magnum, which is a round open area in the middle of the occipital bone. (Figure 1A).
12. Cut the skull open by inserting the lower tip of the small scissors through the foramen magnum and gently cut along the lateral edge of the parietal bone up to the squamosal bone until reaching the eyes (Figure 1A). Use slight pressure to detach the skull from the brain.
13. Using fine-tip forceps, lift the skull from the corner and pull it diagonally up and sideways to expose the brain (see 13). The brain should appear to be a white/pink color without visible blood vessels if the perfusion was successful (Figure 1B-C).
14. Gently remove the whole brain from the skull by detaching the olfactory bulb from the frontonasal suture with the tips of the forceps and cutting the brainstem at the base of the brain (Figure 1A). Place the brain into a 6-well plate filled with ice cold RPMI (see 14).

**Figure 1.**
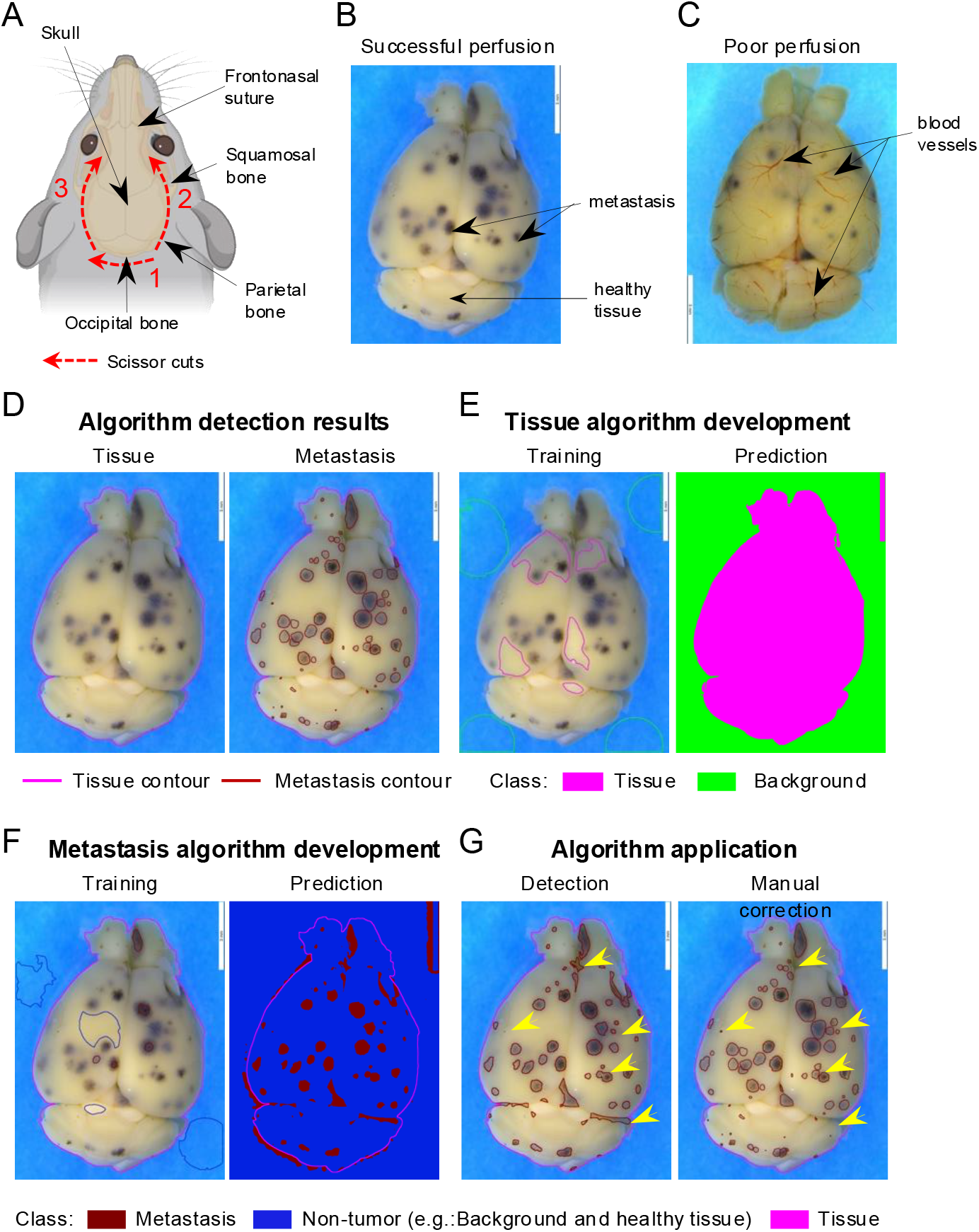
Machine learning algorithm development for brain metastasis quantification. (**A**) Schematic of optimal brain dissection. Red lines depict the incision area, with numbers indicating the order and arrows the direction in which each incision should be made. Dark arrows indicate relevant brain skull regions for the dissection. Figure created with BioRender.com (**B-C**) Representative stereomicroscope images of successfully perfused with no visible blood vessels **(B)** or poorly perfused with visible blood vessels **(C)** brains. Dark arrows indicate representative non-tumor (healthy tissue) brain areas and metastasis (dark dots). (**D-G**) Representative images of the machine learning algorithm development on brain stereomicroscopic pictures using QuPath software (see Note 21). (**D**) Expected brain annotations after running the entire program including tissue detection (left) and metastasis detection algorithm with manual correction (right). Pink line represents the contour of the annotated tissue. Red lines represent the delimitation of each metastasis. (**E**) Tissue detection algorithm development to distinguish brain versus non-tissue areas (Background) as shown in D (Tissue algorithm result picture). Image of representative objects created and manually assigned to a class (“Tissue” or “Background”) by the user during the machine learning training step (left). Automated prediction of the different annotations generated by the software after an optimized training step (right). Brain tissue is represented in pink and the background of the image in green. (**F**) Metastasis detection algorithm development to distinguish metastasis versus non-tumor areas (Background and healthy tissue). Image of representative objects created and manually assigned to a class (“Metastasis” or “Non-tumor”) by the user during the training step (left). Prediction of the different annotations generated by the software after an optimized training step (right). Brain tissue depicted in pink, non-tumor area in dark blue, and metastasis in red. (**G**) Algorithm application. Entire brain annotation obtained after sequentially running the automated tissue and metastasis algorithm (left) and after validating and manually refining (right). Refined annotations are indicated by the yellow arrows.

### 3.3 Imaging

1. Keep all brain samples on ice during imaging. Use RPMI to keep the brain until imaging if cell viability needs to be maintained for further analysis (e.g., flow cytometry) after imaging (see 15).
2. Set up a high-contrast background on the stage of the microscope (see 2).
3. Prepare a petri dish with PBS (∼5 mL) to image the brain and place it on the stage of the microscope.
4. Using the blunt forceps, transfer the brain from the 6-well plate to the PBS petri dish. Make sure that the whole tissue is completely immersed in PBS (see 16 and 17).
5. Using the imaging software from the microscope, image both the dorsal and ventral sides of the brain. Use optimized image acquisition and resolution settings to obtain a clear picture with high definition as shown in Figure 1B. As an example, the settings used for the Leica M165 FC stereomicroscope for our project are described below (see 18 and 19).

a. Image acquisition:

- Ring light intensity: 30
- Transmitted light: OFF
- Exposure time: 52.5 ms (see 19)
- Gain: 0
- Magnification: 4X (objective:0.73 X, eyepiece: 10X, camera adapter: 0.55X)
b. Image resolution:

- Image format: 12.0 Megapixels (4,000 × 3,000), 4:3
- Live format: 2,160 pixels (3,840 × 2,160), 16:9
- Color code: RGB (Red: 32.0, Green: 0, Blue:32.0)
- Color saturation: 50
6. After imaging, the brain can be stored in the desired solution depending on the user’s needs (e.g., formalin, freezing, or RPMI if tissue will be used immediately for further analysis). (see *3* and *15*).

### 3.4 Machine learning algorithm development

#### 3.4.1 Introduction

Our optimized workflow to quantify tumor burden on macroscopic pictures using QuPath software (Bankhead et al., 2017) (see 20, 21, 22, 23, and 24) is organized as below:

1. Creation of a QuPath project and importation of the images of interest (see 25) to initiate the development of the machine learning algorithm and the analysis.
2. Development and validation of two different classifiers (“training” steps) (see 21 and see 26) based on the selection and the classification of structures (“objects”) into two different classes. These steps are performed using the QuPath “Train pixel classifier” function on a group of three or four representative images from the same experiment. It will allow the recognition of:

a. The organ. This algorithm is called “Tissue detection” (e.g., class 1: brain tissue distinct from class 2: the background of the image) (Figure 1D).
b. The metastases. This algorithm is called “Metastasis detection” (e.g., class 1: brain metastasis distinct from class 2: “healthy” brain tissue) (Figure 1D).
3. Automated application of the two algorithms developed on all images (“batch”) from the same experiment to quantify metastatic burden. This results in the complete annotation of all the structures from each image.
4. Manual validation of the tissue and brain metastasis detection.
5. Export and analysis of the numerical data obtained.

#### 3.4.2 QuPath folder and analysis setup

1. Open the QuPath software and create a QuPath project by selecting the command “Create project” in the “Project” tab of the software interface. This will open windows to save this project in the computer workspace (see 21 and *25*)
2. Import three or four representative pictures from the experiment to analyze by selecting them and dragging them into the “Image” tab of the software interface.
3. This will generate a window to indicate the image properties. Apply the settings below:

- Image provider: Default
- Set image type: Brightfield (Other)
- Rotate image: No rotation

#### 3.4.3 Tissue detection algorithm development

1. To create the classes needed for the tissue detection algorithm:

a. Select the “Annotations” tab, then choose the “more options” icon (represented by the three vertical dots in the class window). Click on “Add/Remove” option and select “Add class”
b. Name the new class using the name of the region of interest that needs to be recognized. For instance, here: “Tissue” to recognize brain tissue.
c. Create a second class and name it “Background” as described in 3.4.3.1.a. and 3.4.3.1. b to regroup all regions that are not brain tissue.
2. To initiate the train pixel classifier function, select representative small areas (∼ 0.25 – 1 cm^2^) belonging to each of the two classes and class them manually (Figure 1E, left). This will train the classifier to associate each pixel to one of the two classes through a batch of images (Figure 1D, left). The following steps need to be performed:

a. Select independent pieces of brain tissue and background area using any selection tool from the QuPath toolbar (see 27 and 28).
b. After drawing these objects (see 29), right-click on each object and select Set classification “Tissue” or “Background” according to the nature of the object selected.
3. To train the software to recognize each class through the whole image select “Classify” in the QuPath head menu then select “Pixel classification” and click on “Train pixel classifier”
4. Select the following settings in the pixel classifier window (see 30):

- Classifier: Artificial neural network (ANN-MLP)
- Resolution: Full (352.86 μm/px)
- Features: Default multiscale features ▸ Edit:

∘ 3 channels selected: Red, Green, Blue
∘ Scale: 1.0
∘ Features: Gaussian
∘ Local normalization: None
- Output: Classification
- Region: Everywhere
5. To train and test the classifier click on “Live prediction”. This will apply the training on the whole slide (Figure 1E, right).
6. If the live prediction is not accurate, repeat steps 3.4.3.2 to 3.4.3.5 selecting new areas for each class (see 30).
7. If the algorithm live prediction looks accurate, save the algorithm by entering a name in “Classifier name” section and clicking “save” (e.g., “Tissue Detection”) (see *26*).
8. Test the algorithm on the other representative pictures selected in 3.4.2.2. by selecting “Classify” in the QuPath head menu then “Pixel classification” and clicking on “Load pixel classifier”. Then enter the name of the algorithm on the “Choose model” section and select “Everywhere” as region.
9. If the prediction looks accurate then the algorithm is validated, if not repeat the steps 3.4.3.2 to 3.4.3.5 with a bigger selection of areas across all the images selected in 3.4.2.2 (see 30).

#### 3.4.4 Brain metastasis detection algorithm development

1. Initiate the pixel classifier to recognize metastases and non-tumor areas (e.g., “background” and “healthy brain tissue”) following the same steps as for the tissue detection algorithm described in 3.4.3.1 and 3.4.3.2, but creating two new classes “Metastasis” and “Non-tumor” and select the appropriate areas (Figure 1F, left) (see 31)
2. Train the pixel classifier as described in 3.4.3.3. and 3.4.3.4. but using the following settings (see 30):

- Classifier: Random Trees
- Resolution: Very high (705.72 μm/px)
- Features: Default multiscale features ▸ Edit:

∘ 3 channels selected: Red, Green, Blue
∘ Scale: 2.0, 8.0
∘ Features: Gaussian
∘ Local normalization: Local mean subtraction only
∘ Local normalization scale: 32
- Output: Classification
- Region: Everywhere
3. Test, validate, and save the Metastasis detection algorithm as described from 3.4.3.5 to 3.4.3.9 (Figure 1F, right).

### 3.5 Application of the algorithm

#### 3.5.1 Import batch pictures

1. Create a QuPath project to group all the pictures, annotations, analysis, and measurements as described in 3.4.2.1. (see 32 and 33)
2. Load the images that will be used for the analysis following one of the two options below (see 21):

a. As described in 3.4.2.2.
b. Using the command File ▸ Project▸ Add Images.
3. In both cases, set the image properties as described in 3.4.2.3.

#### 3.5.2 Brain tissue detection on a batch of pictures

1. To apply an existing pixel classifier on a new picture, open the picture and then load the classifier (e.g., here it is “Tissue detection”) by clicking on “Classify” in the QuPath head menu and then selecting ‘Pixel classification” ▸ “Load pixel classifier” (see 34)
2. Select the “Tissue detection” algorithm in the “Choose model” section and “Everywhere” in the “Region” section, this should generate a live prediction as observed during the training classifier step (see 3.4.3.5; Figure 1E, right) (see 35).
3. To create the definitive tissue annotation from the live prediction, the next step is to indicate the size limit expected for the region of interest. This will allow the classifier to define the borders of the annotations. To do so click on “Create objects” tab from the pixel classifier and then select “Full Image” in the “Choose parent objects” section and click “Ok”.
4. Enter the following settings in the “Create Objects” window, (see 36):

- New object type: Annotation
- Minimum object size: 1×10^11^ μm^2^
- Minimum hole size:1×10^12^ μm^2^
- Split objects: selected
5. Confirm the setting by pressing “OK”. This will create and save the final annotation (Figure 1D).
6. As the “Background” annotation is not useful for the rest of the analysis and will slow down the analysis unnecessarily, manually delete this annotation by opening the “Annotations” tab in the left QuPath window and selecting the “Background” annotation and clicking “Delete” (see 37).
7. To automate the analysis, proceed directly to 3.5.3, and then 3.5.4 using the same picture. Otherwise, proceed to the next images by repeating 3.5.2 for each image until the last picture before following 3.5.3.

#### 3.5.3 Brain metastasis detection on a batch of pictures

1. To apply the metastasis detection classifier on the brain tissue, run the respective algorithm in the same way as in 3.5.2. from 3.5.2.1 to 3.5.2.6. but following the modification below:

- Select “Metastasis detection” in the “Choose model” section for 3.5.2.2
- Select “Any annotation ROI” in the “Region” section for 3.5.2.2 (see 38)
- After clicking “Create objects”, select “All annotations” in the “Choose parent objects” section for 3.5.2.3 (see 38).
- For the “create objects” (3.5.2.4.) enter this new parameter size (see 36):

∘ Minimum object size: 1×10^5^ μm^2^
∘ Minimum hole size: 0 μm^2^
∘ Split objects: selected
- Delete the “Non-tumor” annotations for 3.5.2.6 to obtain the final annotation (Figure 1G, left)
2. To automate the analysis, proceed to 3.5.4 using the same picture. Otherwise, proceed to the next images by repeating 3.5.3 for each image until the last picture and then proceed to 3.5.5.

#### 3.5.4 Automatization (see 39)

All the steps described in 3.5. from image setting to both tissue and metastasis detection can be automated by exporting all the commands performed as a script with the user-friendly QuPath interface following the steps below:

1. Upload all the pictures of the project using 3.5.1 without setting the image properties.
2. Run 3.5.1 to 3.5.3 on one picture manually to initialize the script creation.
3. Click Workflow tab (in the left QuPath window), select “Create script”, then click “File” (in the Qupath head menu) and “Save as” to save the script in project file. This will generate a Groovy file in your documents.
4. Click Automate (in the head menu), select “Project scripts” then select the saved script in the menu. This will open the Script Editor window.
5. Click “Run” and the “Run for project”. This will generate “Tissue”, “Background”, “Metastasis”, and “Non-Tumor” annotations on all pictures (Figure 1G, left).
6. In the popup add the image from “available” to “Selected” and click Ok.
7. Delete anything unnecessary (such as “Background” and “Non-Tumor” selections).

#### 3.5.5 Tissue and metastasis annotation accuracy validation and correction

The final annotations generated by the algorithm need to be validated on each picture after the completion of step 3.5.3 or 3.5.4 on all the pictures. Some annotations might need to be rectified (Figure 1G) (see 40). To do so, please follow the steps below:

1. Select the annotation that needs to be corrected in the “Annotations” tab (left QuPath window), then right click select “Unlock”
2. Re-select the same annotation, right click select “Edit single” “Split”
3. Edit the annotation on the picture itself by:

- Refining the annotations border if needed, using the wand tool (can be found in the tool bar) (Figure 1G, right) (see 27)
- Deleting the annotations that are false detection (Figure 1G, right)
- Adding the annotations using the wand tool in the tool bar (Figure 1G, right) (see 21 and 27)

#### 3.5.6 Data export

1. To export the measurements of each annotation generated by the algorithm, select “Measure” (in QuPath head menu) then “Export measurements”. This will generate an “Export measurement” window. These measurements will need to be transformed based on the actual size of the brain (e.g., with Leica M165 FC Stereomicroscope using 1.62 ×10^−10^ to transform BrM area from Qupath measurements to actual size in mm^2^).
2. Select the images of interest for quantification in the “Available” window (right box) and add them to the “Selected” window (right box) by clicking the double right arrow.
3. Fill the tab as below:

- Output file: The desired full path location of the output file
- Export type: The measurement type to be exported (Annotations)
- Separator: comma (.csv)

#### 3.5.7 Data analysis (see 41)

1. Open in excel (.csv) the file generated from QuPath and save it as a (.xlxs) file (see 41).
2. Duplicate the spreadsheet to create a pivot table (spreadsheet 2) without affecting the original data (spreadsheet 1). The pivot table will allow you to organize the data output in a meaningful way.
3. Highlight the first row, select “Data” then “Filter” (see 42)
4. Insert a new column named “Side” between column A (Image) and B (Name) containing which side of the brain is visible on the picture (e.g., dorsal or ventral).
5. Insert another column after column A labeled “Sample ID”.
6. Change the “number” format in column J (Area -µm^2^) to “scientific” format
7. Select whole table then select “Insert” and click “Pivot Table” then “New Worksheet”. This will generate the “PivotTable Fields” window.
8. Select “Group”, “Sample ID, “Side”, “Class” (QuPath annotation name column, e.g., “tumor”, “tissue”), and “Area -µm^2^”, and drag them from “Field Name” to “Rows”.
9. Drag “Name” from “Field Name” section to “Filters”.
10. Drag “Count of Class” and “Sum of Area -µm^2^” to “Values”.
11. The result should be a pivot table organized by experimental group indicating the number of metastatic lesions and the total metastatic area for each side of each brain (see 41).
12. Plot the total number of metastases, total metastatic area per mouse, and individual metastatic lesion size per group. It is recommended for the first time to verify the accuracy of the method by comparing the number of metastases count manually on the pictures and the count provided by the algorithm classifier run (Figure 2 A-E).

**Figure 2.**
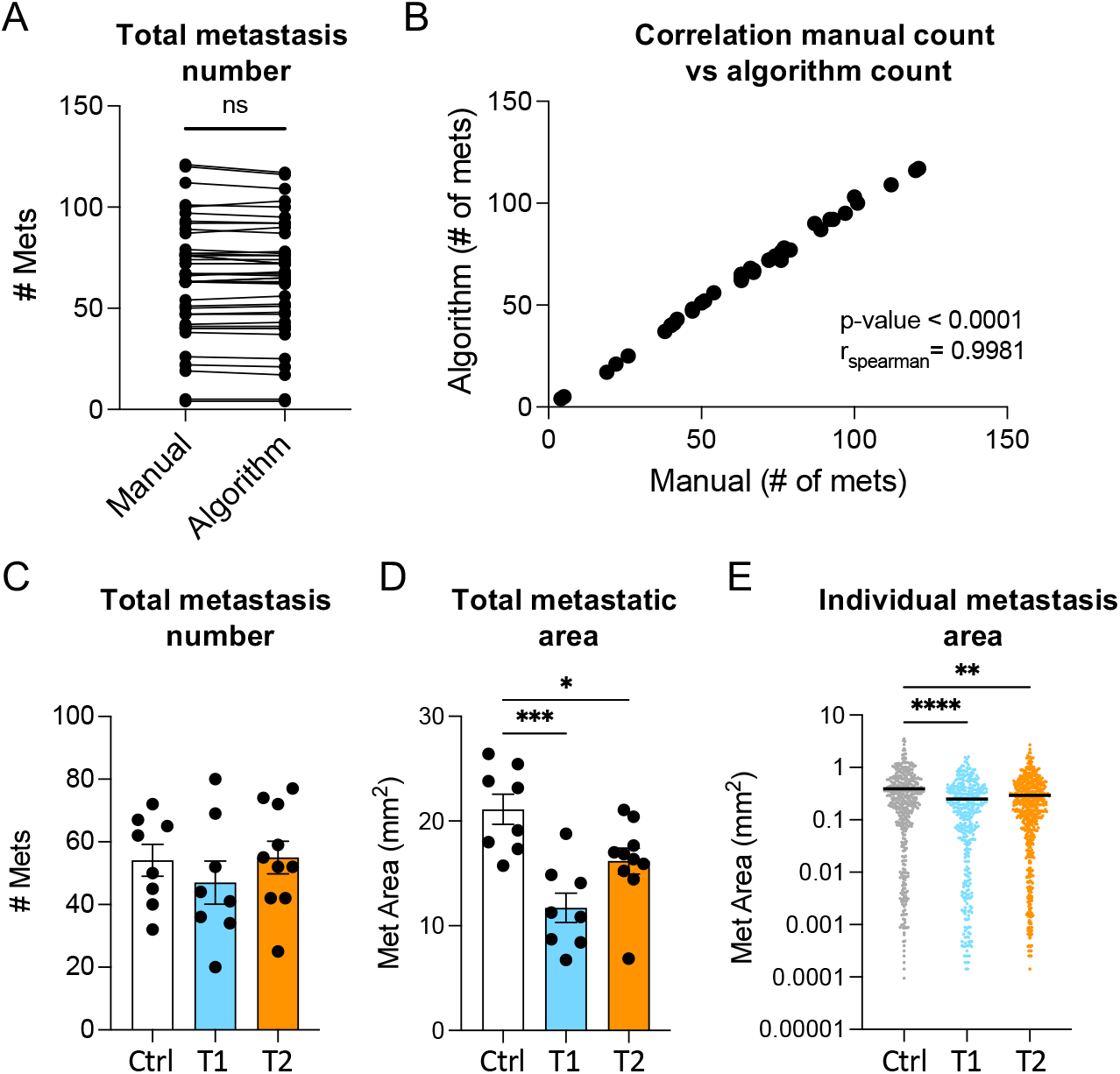
Analysis of brain metastatic (BrM) burden applying the machine learning approach. (**A-B**) Representative comparison of BrM number obtained by manual count (Manual) or automated count (Algorithm). (**A**) BrM count. (**B**) Spearman correlation. (**C-D**) Representative data obtained with the machine-learning approach comparing different experimental groups [control (Ctrl), treatment 1 (T1), and treatment 2 (T2)]. (C) Total number of brain metastases per mouse in each group. (D) Total metastatic area per mouse in each group. (E) Size of each individual metastasis per group. Data in A compared using paired t-test. Data in C-D compared using one-way ANOVA followed by Tukey correction. Data in E compared using Kruskal-Wallis test followed by Dunn’s correction. *p < 0.05, **p < 0.01, ***p < 0.001, ****p < 0.0001.

## 4 Results and discussion

Here, we developed an approach to accurately quantify metastatic burden on macroscopic organ images using the open-source, user-friendly software QuPath. We describe the development and application of machine learning algorithms based on pixel classification to detect tumoral and healthy areas in the tissue (Figure 1G). We demonstrated that the developed classifiers and workflow enable the accurate detection and precise delimitation of all brain metastatic lesions (Figure 1D-G). When comparing BrM quantification using our method with manual counts from multiple samples, we found a similar BrM number and a strong correlation between the two approaches, thus validating the accuracy of our algorithms (Figure 2A-B). However, unlike manual counts (Bankhead et al., 2017; Lopès et al., 2018; Pennig et al., 2021), our automated pipeline offers a more time-efficient approach. These classifiers are exportable and can be easily shared between users, thereby, minimizing user bias and increasing reproducibility across experiments.

We used our approach to evaluate BrM burden in three different experimental groups (Figure 2C-E). Although we did not observe a difference in BrM counts, we found a decrease in total metastatic area and individual lesion size among groups (Figure 2C-E). Therefore, our method revealed variations that may have not been captured by traditional counting. Importantly, this method gives access to data points not always accessible by other techniques, such as individual metastatic lesion size (Figure 2E). This BrM burden indicator is crucial to consider when testing therapeutic agents for brain tumors, as BrM size not only influences patient prognosis but also guides the selection of the most appropriate treatment regimen (Amsbaugh and Kim, 2024; Li et al., 2022; Santos et al., 2024). This underscores the significance of the tool we are providing for preclinical studies.

While we describe the applicability of the method on brain metastases, it can also be adapted to quantify tumor burden in other models and organs. An important advantage of this approach is that it preserves the tissue integrity after imaging, allowing for further characterization of the same sample, such as immune profiling, transcriptomic and/or metabolic analyses. However, it does come with limitations. For example, because it requires organ harvest, it is not suitable for longitudinal studies. Additionally, lesions need to be distinguishable from healthy tissue (e.g., melanocytic lesions or lesions with a recognizable color or shape), although this can be addressed by using a reporter protein (see 22), albeit with the introduction of other drawbacks (see Table 1). Notably, if a fluorescent reporter protein is used, the tumor detection algorithm needs to be adapted for fluorescence settings, although the overall workflow and tissue detection remain the same.

This protocol can measure all superficial lesions, but deeper lesion assessment requires cutting the organ at different depths. While this introduces some limitations similar to the ones described for histology (Table 1), it will still preserve the tissue for further use (see 23). Additionally, this protocol can be extended to relatively flat primary tumors; however, it is not recommended for models such as subcutaneous tumors as it would require taking multiple pictures to assess the total tumor volume and it will be more time-consuming and less accurate than the traditional caliper measurement. Nevertheless, this method represents a valuable tool for accurately assessing metastatic burden while allowing the correlation of relevant biological variables for the study of disease progression, the tumor microenvironment, and response to therapy.

## 5 Method and troubleshooting notes

1. Product catalog numbers and sources are provided as a reference but equivalent products from other sources can be used.
2. The use of a colored background during imaging helps the distinction between brain tissue and background by the algorithm. This is optional but recommended as it will gain a significant amount of time for the algorithm setup. After multiple tests, we determined that the use of a blue background was the most efficient. A post-it placed under the petri dish containing the brain organ can be used.
3. After imaging, the brain can be used further analysis (e.g., flow cytometry) or stored for future use. The brain tissue should be placed in the appropriate buffer immediately after imaging and this buffer should be prepared in advance. For instance: 10 mL of cold RPMI for flow cytometry or 10% Formalin (Sigma, catalog: HT501129-4L) for histology.
4. The best forceps style for brain harvest is curved to allow optimal extraction of the whole brain organ from the skull. Blunt forceps are the best to manipulate the brain during the imaging step without damaging the tissue.
5. Other microscopes with similar features can be used. The settings depend on the equipment, updates of the equipment and software, and the tissue, therefore the imaging settings written in this protocol are indicative and will likely need to be adjusted to fit specific user equipment.
6. Using a ring light as the light source for the microscope provides the best images, but it is not mandatory.
7. Microscope software will depend on the brand of the microscope.
8. Other injection method (e.g., intracranial or intracardiac) can be used as well. In the case of implantable tumor models, the number of cells will vary depending on the model and appropriate institutional ACUC guidelines. The harvest time point should be determined depending on the model being used.
9. The syringe should not contain any air bubbles to allow the best perfusion and avoid damaging organs and blood vessels.
10. To avoid blood clots and preserve the brain tissue, it is recommended to harvest one or two mice per person at a time.
11. For more information and illustrations, a detailed perfusion protocol can be found in the literature (Wu et al., 2021).
12. This step needs to be performed carefully, making sure to not cut the heart or any major blood vessels before perfusion.
13. To help lift the skull without damaging the brain, it is suggested to cut the skull along its middle line.
14. The brain tissue is very fragile and needs to be harvested very carefully to avoid damage.
15. Brains should be imaged in batches directly after harvesting within the same time and with the same settings. The brain should not be left in PBS or RPMI for extended periods of time as it will begin to inflate. After fixation, the brain will shrink distorting its shape. Thus, it is important to handle brains from a given experiment in the same manner to avoid technical bias.
16. To prevent drying, the brain must be fully covered with PBS. Avoid overflow and air bubbles for picture quality purposes. A transfer pipette can be used to add and remove PBS as well as air bubbles.
17. The Petri dish needs to be changed regularly to avoid debris in images. If the brains are used after imaging for other analysis, the Petri dish needs to be changed between each brain to avoid contamination.
18. For each microscope/software brand and serial number, the settings for imaging might need to be adjusted.
19. If the brain does not have a lot of metastatic lesions, a lower exposure is recommended.
20. MAC, Windows or Linux can be used to run QuPath. The minimum requirement is a 64 bits environment. An intel core i7 or above hardware with 16GB RAM is recommended.
21. A detailed description of the QuPath software and tutorials are available online at: https://qupath.github.io/
22. This method works with pigmented or visually detectable lesions, another approach would need to be used to visualize unpigmented lesions. Black ink can be used to make the metastasis visible. Alternatively, a xenobiotic marker can be used to visualize and image the metastasis with an appropriate fluorescent microscope. Metastasis burden can then be quantified on the images using our method by adapting it for fluorescent imaging. The latter might artificially increase tumor immunogenicity (Table 1).
23. This method works for metastases visible on the surface of the organ. To apply it on deeper metastases it is required to slice the organ at the different depths needed and image each tissue section. In this case all tissue section should be keep in cold RPMI media until other use of the tissue.
24. This protocol describes the use for brain metastasis but can also by applied on other organs.
25. This project will contain all pictures and annotations used by the newly developed algorithms. The new project folder must be empty before use and created only for QuPath analysis to avoid any data and analysis loss.
26. The classifiers developed are then saved in a file and can be loaded and used in any QuPath software. They can be shared between users.
27. The “wand tool”, which is the 9^th^ icon from Qupath tool bar, is the best selection method as it is based on pixel value and not only user criteria. It is thus recommended to use this tool, but any others would still work.
28. It is crucial to have only one type of class/tissue in each selection. Area of interest should not be mixed (e.g., tissue and background selected objects should respectively contain only tissue or background).
29. The number of objects selected depends on the complexity of the tissue but an optimal 5 to 8 objects by class is preferable.
30. These settings can vary according to imaging equipment and tissue complexity. Thus, these might need to be adjusted for optimal results. As a general tip, artificial neural network classifiers are better for discriminating uniform area with clear differences (such as background versus tissue), while random trees classifier are more helpful to distinguish area with more finer differences and harboring a lot of pixel color and intensity diversity. Highest resolution will perform better in detecting detail and small objects (such as metastasis) while lowest resolutions are the best to detect global structure (such as tissue). The features option is relevant to use when RGB color differences are observed between class and might need to be tested to find out the optimal combination of settings.
31. The previously recommended size of the area selected needs to be adapted for the metastatic lesion. It is recommended to do multiple selections of different lesions by selecting each entire chosen lesion.
32. All pictures need to be analyzed using the same algorithm within the project.
33. Tif images as well as Aperio (.svs, .tif), Hamamatsu (vms, .vmu, .ndpi), Leica (.scn), MIRAX (.mrxs), Philips (.tiff), Sakura (.svslide), Trestle (.tif), Ventana (.bif, .tif), Generic tiled TIFF (.tif), ImageJ TIFF, JPEG, PNG are compatible with QuPath.
34. This can be used to apply the classifier to new images, but not to continue training.
35. If the live prediction does not accurately represent the different annotations, it is preferable to not pursue and improve the algorithm before applying it.
36. These settings are optimal for the model used as an example in this protocol. They can be used as a starting point for any other model or tissue but will likely need to be adjusted. Notably, the size of the object will depend on the size of the tissue being imaged.
37. It is preferable to delete any annotations that will not be used for further analysis to decrease the time of analysis and to avoid any confusion between unnecessary area.
38. With these settings the algorithm only runs within the tissue detection.
39. Make sure to have run the tissue detection and metastasis detection on one picture before to do this step. It will not work otherwise.
40. In general, the tissue detection does not need any correction. In contrast, it is more frequent that the metastasis detection requires correction. It is recommended to improve the algorithm if more than 20% of each image needs to be corrected.
41. Make sure to check and validate all tissue and metastasis annotations before proceeding with this step.
42. Select tumor from the filter on top to make sure you are only looking at tumor measurements, not tissue.

## Acknowledgements

We thank April Huang and other members of the Goldszmid laboratory for their technical help and feedback. We gratefully acknowledge our collaboration with Dr. Merlino (CCR-NCI) in developing the metastatic model used as an example in this protocol. We also thank him for providing access to the LEICA microscope and software. This research was supported by the Intramural Research Program of the NIH (CCR-NCI) and a CCR FLEX Synergy Award.

